# Comparative Molecular, Innate, and Adaptive Impacts of Chemically Diverse STING Agonists

**DOI:** 10.1101/2025.02.21.639458

**Authors:** Nobuyo Mizuno, Dylan Boehm, Kevin Jimenez-Perez, Jinu Abraham, Laura Springgay, Ian Rose, Victor R. DeFilippis

## Abstract

Pharmacologic activation of the innate immune response is being actively being pursued for numerous clinical purposes including enhancement of vaccine potency and potentiation of anti-cancer immunotherapy. Pattern recognition receptors (PRRs) represent especially useful targets for these efforts as their engagement by agonists can trigger signaling pathways that associate with phenotypes desirable for specific immune outcomes. Stimulator of interferon genes (STING) is an ER-resident PRR reactive to cyclic dinucleotides such as those synthesized endogenously in response to cytosolic dsDNA. STING activation leads to transient generation of type I interferon (IFN-I) and proinflammatory responses that augment immunologically relevant effects including antiviral responses, antigen presentation, immune cell trafficking, and immunogenic cell death. In recent years engineered cyclic dinucleotides and small molecules have been discovered that induce STING and safely confer clinically useful outcomes in animal models such as adjuvanticity of anti-microbial vaccines and tumor clearance. Unfortunately, clinical trials examining the efficacy of STING agonists have thus far failed to satisfactorily recapitulate these positive outcomes and this has prevented their translational advancement. A likely relevant yet perplexingly under investigated aspect of pharmacologic STING activation is the diversity of molecular and immune responses that associate with chemical properties of the agonist. Based on this, a comparative survey of these was undertaken using unrelated STING-activating molecules to characterize the molecular, innate, cellular, and immune outcomes they elicit. This was done to inform and direct future studies aimed at designing and selecting agonists appropriate for desired clinical goals. This revealed demonstrable differences between the agonists in potency, transcriptomes, cytokine secretion profiles, immune cell trafficking, and antigen-directed humoral and cell mediated immune responses. As such, this work illustrates that phenotypes deriving from activation of a protein target can be linked to chemical properties of the engaging agonist and thus heightened scrutiny is necessary when selecting molecules to generate specific *in vivo* effects.

## INTRODUCTION

Subunit vaccine antigens when used alone do not typically exhibit levels of immunogenicity that are sufficient for the establishment of protective immunity. As such, they require coadministration of factors such as adjuvants to stimulate the innate immune system in a manner resembling that which occurs during exposure to molecular signals of proliferative microbial danger [1]. Unfortunately, only a small number of adjuvants are approved for clinical use and effective protection requires pathogen-specific immune polarization such that one adjuvant is unlikely to achieve all objectives [2]. Moreover, the precise mechanisms of action of clinical and experimental vaccine adjuvants are often poorly characterized [3]. The need is therefore great for pursuit and characterization of new adjuvant classes, especially due to the likelihood of pathogen emergence and the number of established infections for which no or suboptimal vaccines exist. A promising investigative approach to adjuvant discovery is identification of pharmacologic activators of pattern recognition receptors (PRRs) [4]. Engagement of PRRs by pathogen- or danger-associated molecular patterns initiate intracellular signaling processes that culminate in innate immune activation. This includes proinflammatory cytokine secretion and activation of antigen presenting cells (APC) that facilitate and direct antigen-directed adaptive immune responses.

<u>St</u>imulator of <u>in</u>terferon <u>g</u>enes (STING) is an ER-resident PRR reactive to cyclic purine dinucleotides (CDNs) synthesized by bacterial and eukaryotic nucleotidyl cyclases. Most prominently, cyclic GMP-AMP (cGAMP) synthase (cGAS) is a vertebrate enzyme activated following contact with cytosolic DNA to synthesize 2’3’ cGAMP from ATP and GTP. Activated STING leads to phosphorylation of TANK binding kinase 1 (TBK1) which in turn phosphorylates transcription factors such as interferon regulatory factor 3 (IRF3) and nuclear factor κB (NF-κB). These leads to expression of type I interferons (IFN-I) as well as other pro-inflammatory cytokines and costimulatory factors. In addition, STING is expressed across a wide range of cell types and induces transcription-independent processes such as autophagy, inflammasome induction, and lysosomal biogenesis often in ways that are cell- and agonist-specific [5–8]. These collectively contribute to the innate immune phenotypes conferred by activated STING and facilitate the establishment of adaptive immune responses through localized recruitment and maturation of APCs, antigen loading and processing, and antigen presentation in the draining lymph node (dLN). STING is therefore an increasingly attractive cellular target for development of new adjuvants and its stimulation as such has shown to enhance efficacy of vaccines against diverse pathogen types.

Multiple CDN classes can activate STING and enhance vaccine-mediated immune responses in animal models. Furthermore, synthetic small molecules have also been identified that induce STING and adjuvant functions have been described for many of these. Numerous efforts are currently underway to thus discover and characterize novel STING-activating molecules for similar use. However, liabilities associated with certain chemical classes can affect their *in vivo* suitability, potency, and performance. For instance, CDNs are susceptible to degradation by phosphodiesterases and also cross cell membranes poorly due to their size and hydrophilicity. More importantly, the profile of STING-mediated molecular processes is influenced by the chemical structure of the ligand. This may be due to a number of causes including engagement of off-target proteins, pharmacokinetics, region of binding, or molecule-specific conformational changes in STING that affect its activity. As such, it is likely insufficient to simply target STING using any activating ligand since the consequent immune phenotypes could vary substantially and confer outcomes inconsistent with a vaccine’s purpose.

This study aims to undertake *in vitro* and *in vivo* characterization of three well described but chemically dissimilar STING agonists. Our purpose is to perform direct comparisons of molecular, innate, and adaptive immune responses that they induce to identify effects that are both shared and divergent between them. We hope these will inform future studies that pursue STING inducers for specifically defined clinical outcomes. This work also highlights multi-level phenotype patterns correlating with STING activation that were unexpected in our models.

## RESULTS

### Dose-dependent activity of STING agonists in vitro

To compare molecular, innate, and adaptive immune phenotypes elicited by chemically dissimilar STING ligands we used cyclic di-AMP analog ML-RR-S2 CDA (CDA) [9], the amidobenzimidazole diABZI [10], and 5,6-dimethylxantheonone-4-acetic acid (DMXAA) [9,11,12]. We first characterized the potency, patterns of innate activation, and cytotoxicity to allow identification of suitable doses for subsequent *in vivo* use. For this we used RAW264.7 and J774 macrophage-like murine cell lines engineered to express luciferase in response to IFN-I-dependent signaling [13,14]. These were exposed in duplicate to a range of concentrations of the molecules and luciferase signal measured after 24 h. CDA was transfected since it is not easily cell permeable due to its size and hydrophilicity. In parallel, cytotoxicity was also measured using the ATP-based Cell Titer Glo reagent. As shown in **Figures 1A and 1B**, differences were observed between the cell types primarily at the highest concentrations of each molecule with RAW264.7 displaying overall lower viability than J774 cells. Moreover, J774 cells also showed greater sensitivity to the compounds as indicated by higher overall fold changes in luciferase induction for all agonists. Additionally, CDA elicited the highest peak induction levels in both cell types. J774 cells co-express secreted alkaline phosphatase (SEAP) in response to NF-kB activation and we therefore also examined this response. As shown in **Figure 1C**, detectable SEAP was observed at the highest dose of DMXAA and at multiple doses of CDA but was not detected at any diABZI dose. Importantly, these results indicate that levels of cytotoxicity at doses eliciting peak innate induction in both cell types were not unsuitably low indicating that these were appropriate for use in additional studies. Moreover, NF-kB activation was only significantly detected in response to CDA, highlighting a key difference in phenotypic response to the agonists.

**Figure 1.**
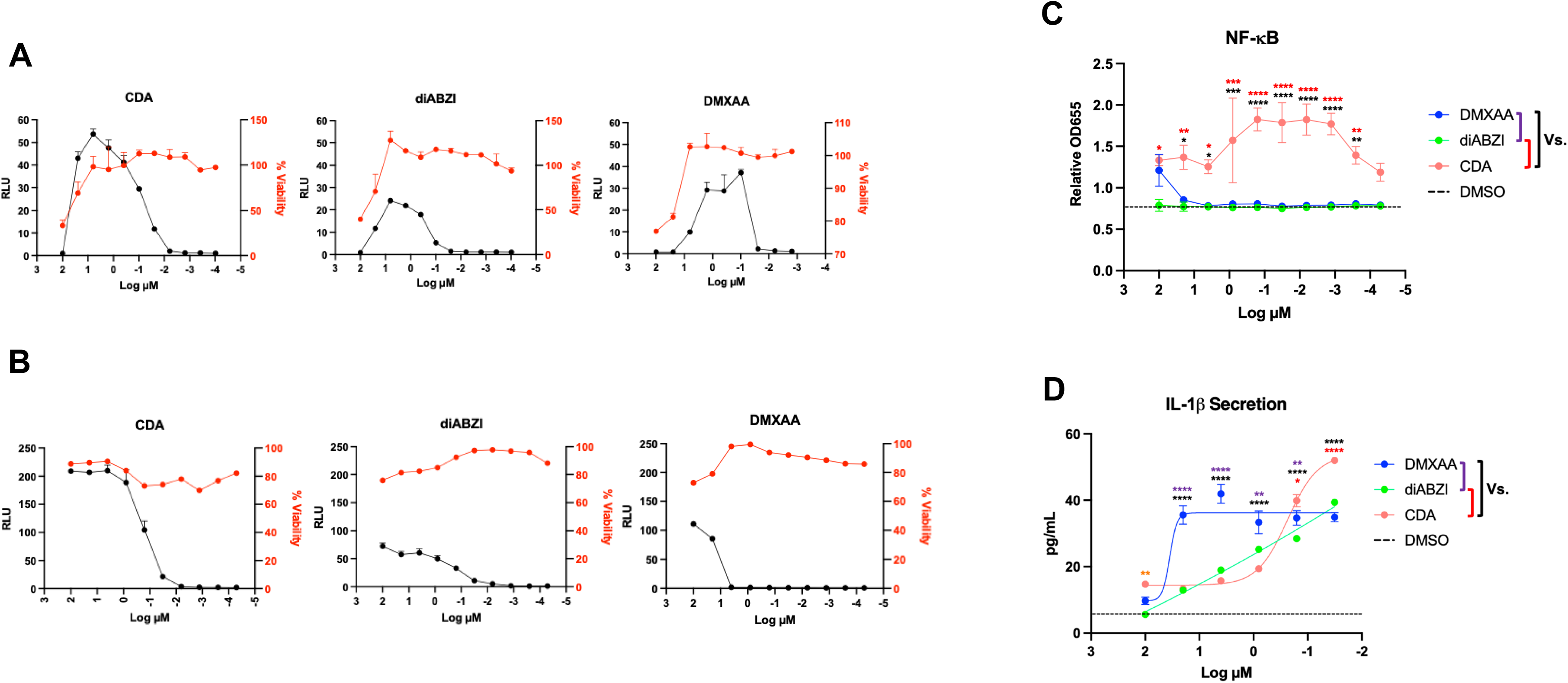
Dose-dependent innate responses induced by STING agonists in murine cell lines. RAW264.7 (A) and J774 (B, C, D) were grown to confluence in 96 well plates and exposed in duplicate overnight to CDA, DMXAA, or diABZI at indicated molarities. A. Cells were exposed to either Quanti-Luc or Cell Titer Glo lysis buffer to measure IFN-I signaling induction or cell viability, respectively. Data presented are mean ± SEM relative luminescence units (RLU, black) or percent viability (red) determined relative to control cells treated with DMSO- or transfection reagent (CDA); C. NF-kB-dependent SEAP expression expressed as mean ± SEM absorbance at OD655 for indicated treatment relative to control treated cells; D. Secretion of IL-1b from primed J774 cells treated with indicated doses of DMXAA, diABZI, or CDA. Data displayed are mean ± SEM IL-1b pg/mL secreted into culture media. Two-way ANOVA with Tukey’s correction for multiple comparisons was used to examine statistical significance of dose-specific activation of NF-kB or IL-1b secretion between agonists (*p < 0.05, **p < 0.01, ***p < 0.001, ****p < 0.0001 color coded as indicated for agonist pairing).

Activation of STING is also known to stimulate the NLRP3 inflammasome, which can greatly affect the proinflammatory tissue environment through secretion of IL-1β and IL-18 [5,15]. However, the degrees to which this is induced by different agonist chemical classes has not been examined. We therefore used the J774 monocyte cell line as has been previously described for NLRP3 inflammasome studies to assess the response to these agonists [16,17]. In duplicate cells were primed for 5 h with LPS and exposed overnight to a dosing range of diABZI, CDA, or DMXAA. Secretion of IL-1**β** was then measured using bead-based Luminex assay. As shown in **Figure 1D**, all three agonists stimulated secretion of IL-1β although with divergent induction patterns that were maximal near 0.032 µM. However, while diABZI secretion declined linearly with higher agonist doses, DMXAA and CDA generated dissimilarly sigmoidal decreases with higher dosage. Based on these results we conclude that the agonists collectively stimulate qualitatively different patterns of intracellular innate induction as indicated by degree of dose-dependence, cytokine (IFN-I and IL-1β) secretion, and NF-κB activation.

### STING agonist-mediated dendritic cell maturation

Adjuvant-enhanced adaptive immune processes require the local activation of dendritic cells (DC) to enable antigen uptake and presentation on MHC-II to CD4^+^ helper cells in the draining lymph node (dLN) [18]. We therefore compared the ability of STING ligands to induce the expression of MHC-II and costimulatory proteins CD86, CD80, and CD40. For this we used nontoxic concentrations of each molecule near those that displayed the highest reporter signal in *in vitro* assays (**Figure 1**). Control stimuli included LPS and Alum as respective inducers of DC activation and subunit vaccine response enhancement. Immature bone marrow derived DC (BMDC) from C57Bl/6 mice were treated overnight and flow cytometry used to measure surface expression of the markers. As shown in **Figure 2**, all STING ligands as well as LPS led to significant upregulation of the four markers relative to DMSO-treated cells whereas no such induction was observed for Alum. In addition, DMXAA appears capable of eliciting significantly higher levels of expression of these proteins than the other STING ligands and (with the exception of MHC-II) LPS. As such, these STING agonists are highly but differentially effective at inducing maturation of DC in an *ex vivo* system, a process necessary for establishing adaptive immune responses.

**Figure 2.**
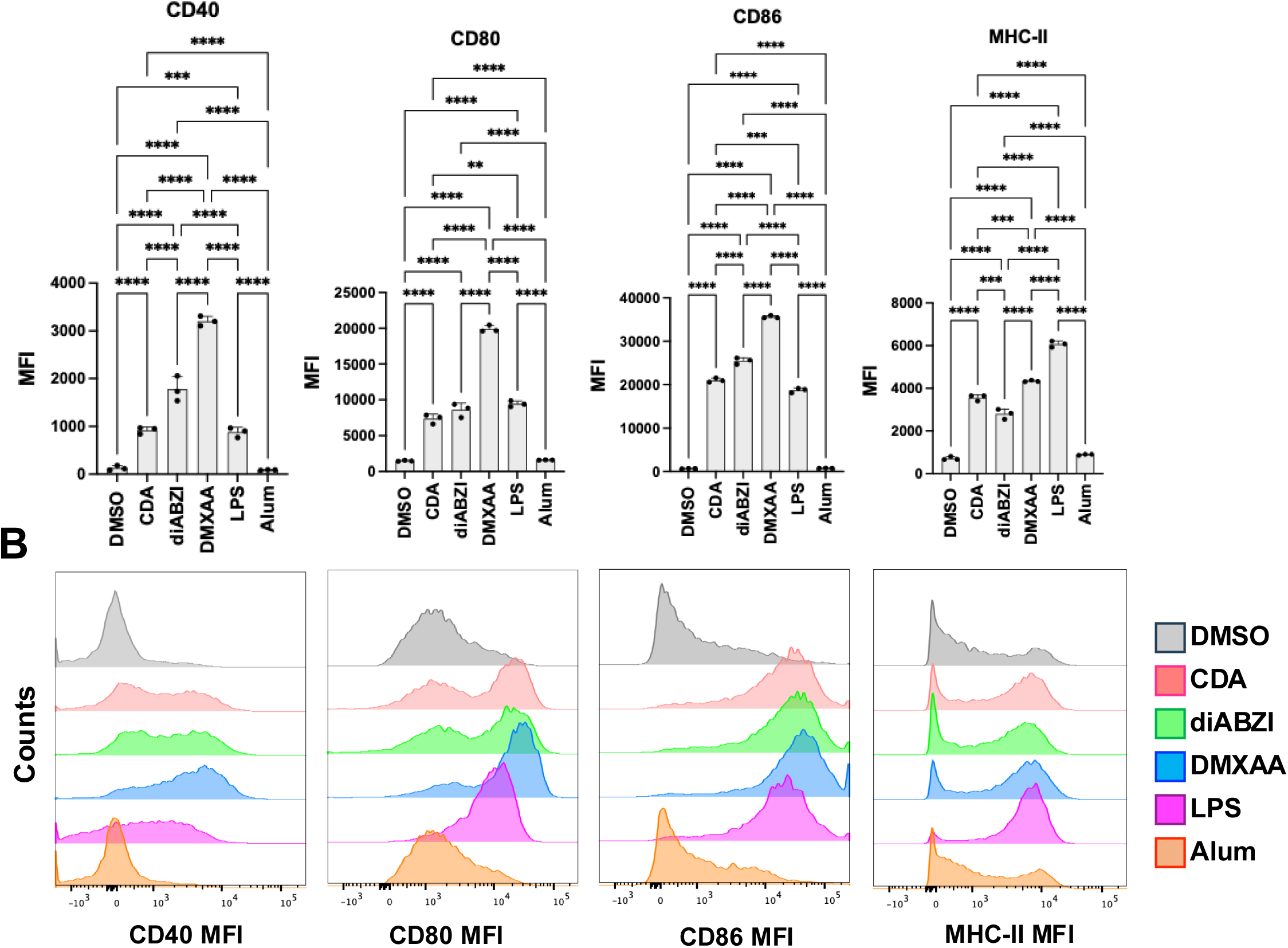
Maturation of Dendritic Cells from C57Bl/6 Mice in Response to Innate Stimuli. Immature bone marrow derived DC (BMDC) from C57Bl/6 mice were treated with 0.5% DMSO, 1 µM diABZI, 50 μM DMXAA, 1 µM CDA, 100 ng/mL LPS, or 3 µg/mL Alum as indicated. Cells were collected after 24 h and stained for expression of CD40, CD80, CD86 and MHC-II and analyzed by flow cytometry. **A.** Mean fluorescence intensities (MFI) ± SD for indicated treatment and surface marker. One way ANOVA with Tukey’s multiple comparisons test was used to compare results between treatments (***p < 0.001****; p < 0.0001); **B.** Representative MFI histograms for each marker are shown for cells exposed to indicated stimulus.

### In vivo cytokine induction by STING agonists

Adjuvant mechanisms associate with stimulation of innate processes immediately following *in vivo* administration. While administered locally, this often manifests as secretion of proinflammatory cytokines that are detectable in peripheral blood [19,20]. We therefore compared this indicator of systemic innate immune induction by the three molecules using secretion of peripheral cytokines as a readout. Using doses that are consistent with what was found to induce near maximal IFN-I levels, we injected each compound intramuscularly (IM) into C57Bl/6 mice to simulate a typical vaccination route using PBS as a negative control injection. Mice were euthanized at 24 h and peripheral blood collected. Levels of cyto/chemokines were then measured in serum using Luminex multiplex bead-based assay. As shown in **Figure 3**, three different cytokine induction patterns were detected for the stimuli. Secretion of cytokines such as CCL7 (MCP-3) and IL-1β was significantly and similarly induced by all three treatments relative to control mice. Cytokines such as CXCL10 and IL-18 were significantly stimulated by CDA and DMXAA but not diABZI. Finally, DMXAA alone induced significant secretion of CCL2, CCL3, CCL4, CCL5, CCL11, CXCL1, CXCL2, IL-6, IL-10, IL-12p70, IL-17A, IL-22, IL-27, and TNFα. These results suggest that while some innate pathways such as those leading to secretion of CCL7 and IL-1β are stimulated comparably in response to all stimuli, others can be differentially activated in an agonist-specific manner. Moreover, the response to DMXAA appears unusual among the agonists in that it is uniquely capable of inducing secretion of a subset of cytokines. This is underscored by the fact that no secretion of CCL3, IL-10, IL-17A, IL-22, and IL-27 above the limit of detection was observed in response to other agonists for yet they were significantly stimulated by DMXAA.

**Figure 3.**
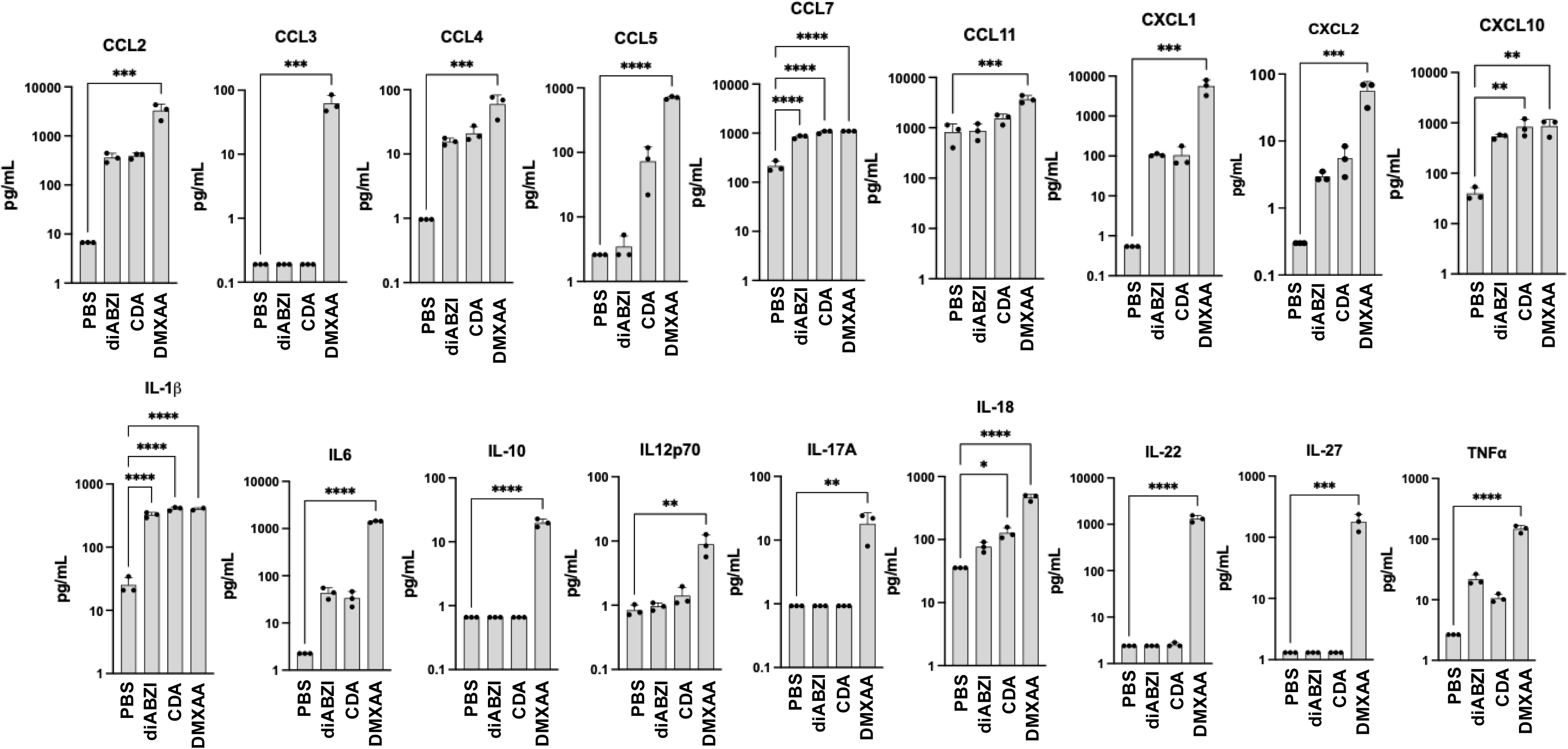
Induction of systemic cytokine secretion by STING agonists *in vivo*. C57Bl/6 mice were injected IM with PBS, 30 µg diABZI, 10 µg, CDA, or 500 µg DMXAA and serum harvested at 24 h. Luminex bead based multiplex assay was then used to measure absolute levels of indicated cytokines. Data presented are mean ± SD pg/mL. One way ANOVA with Dunnet correction for multiple comparisons was used to determine statistical significance (**p <0.01, ***p < 0.001, ***p < 0.0001).

### In vivo transcriptional impact of STING agonist administration

The rapid innate processes conventionally induced by adjuvants are most highly localized to the site of administration and dLN where adaptive immune processes commence. We next examined this at the transcriptional level by injecting mice with diABZI, CDA, DMXAA, or PBS as described above. Total RNA was isolated from dLN harvested at 5 h, 24 h, or 72 h and semi-quantitative RT-PCR (qPCR) used to measure adjuvant-dependent upregulation of mRNAs encoding conventional IFN-stimulated and pro-inflammatory proteins. As shown in **Figure 4A**, transcriptional induction peaked for nearly all genes at 5 h and declined in later time points although at different rates between agonists. Importantly, levels of all examined mRNAs displayed significant differences between at least two stimuli at multiple time points with the most differences observed between DMXAA and CDA and the fewest between DMXAA and diABZI. Moreover, DMXAA stimulation led to the highest mean induction of all genes.

**Figure 4.**
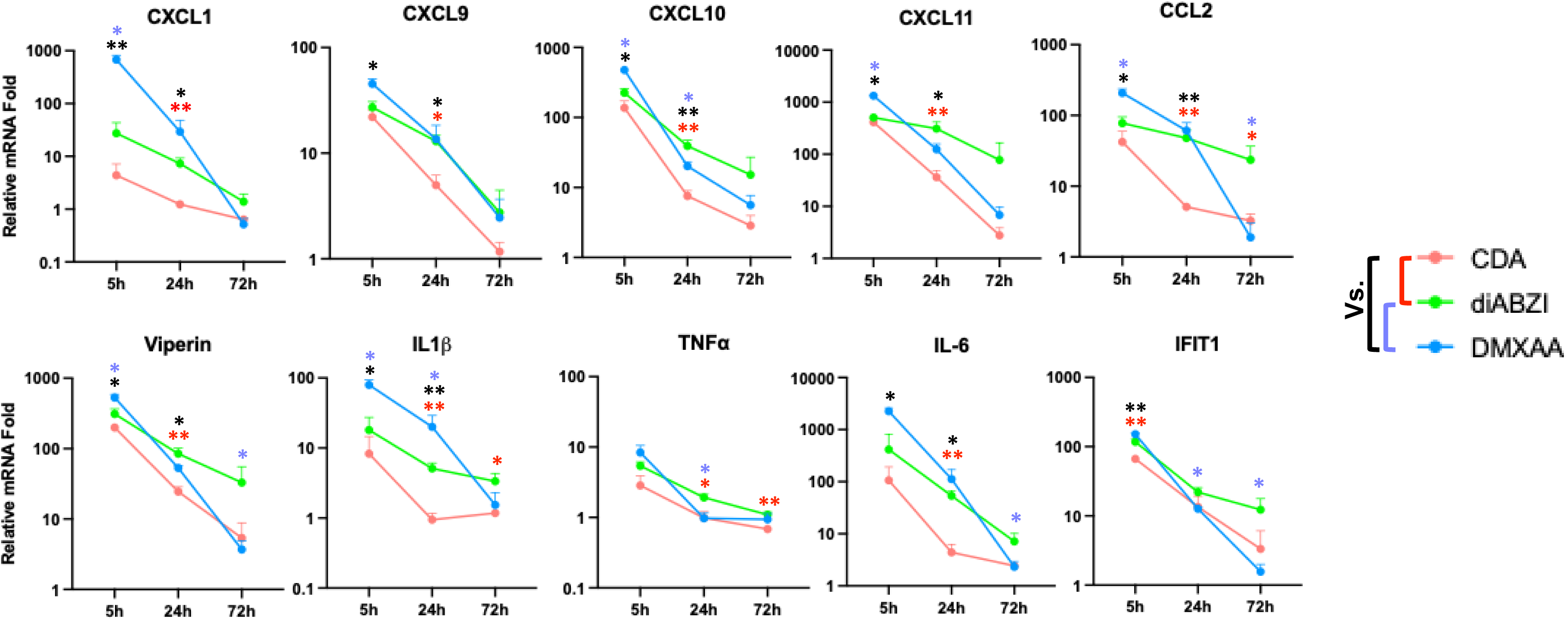
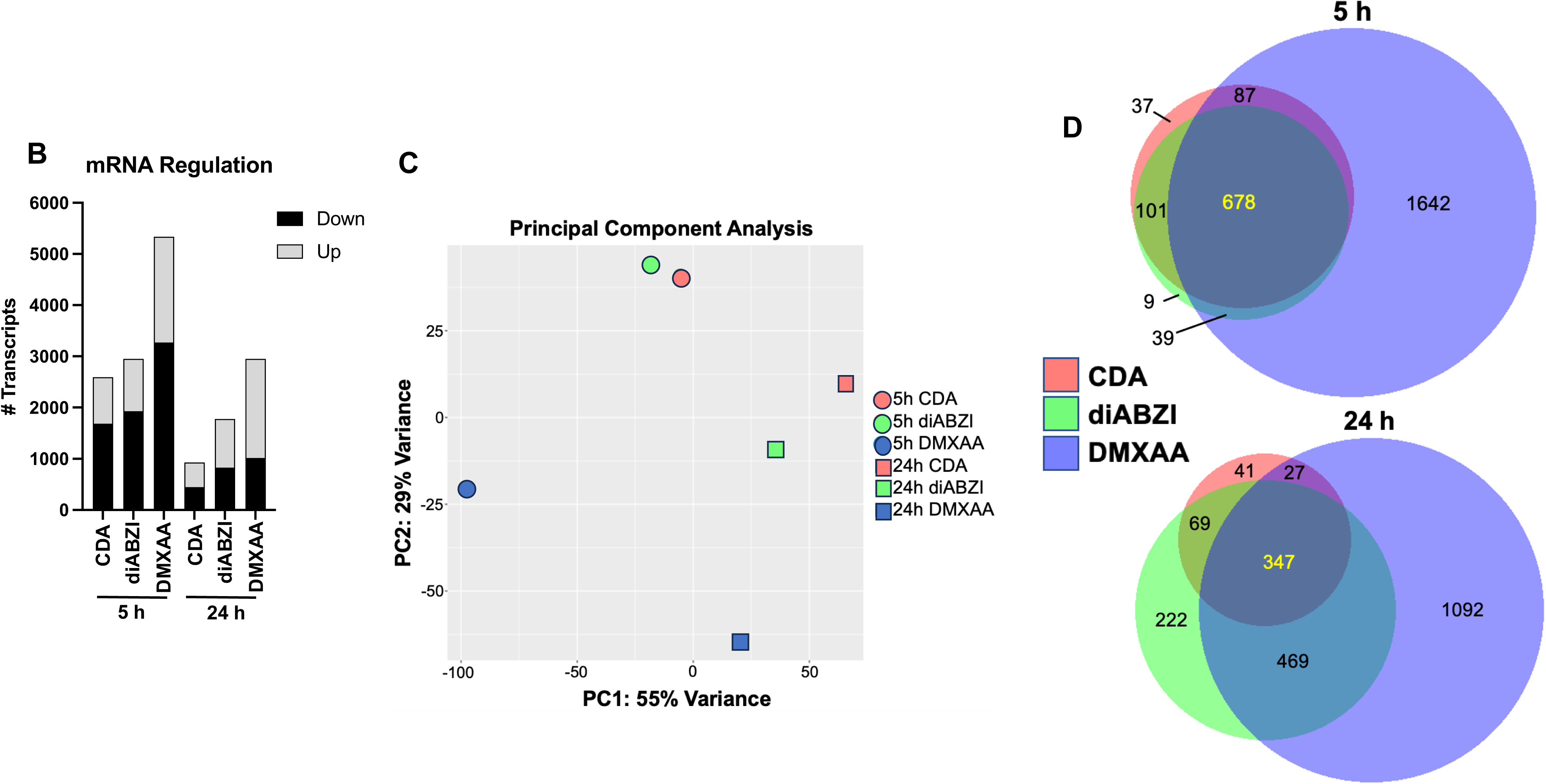
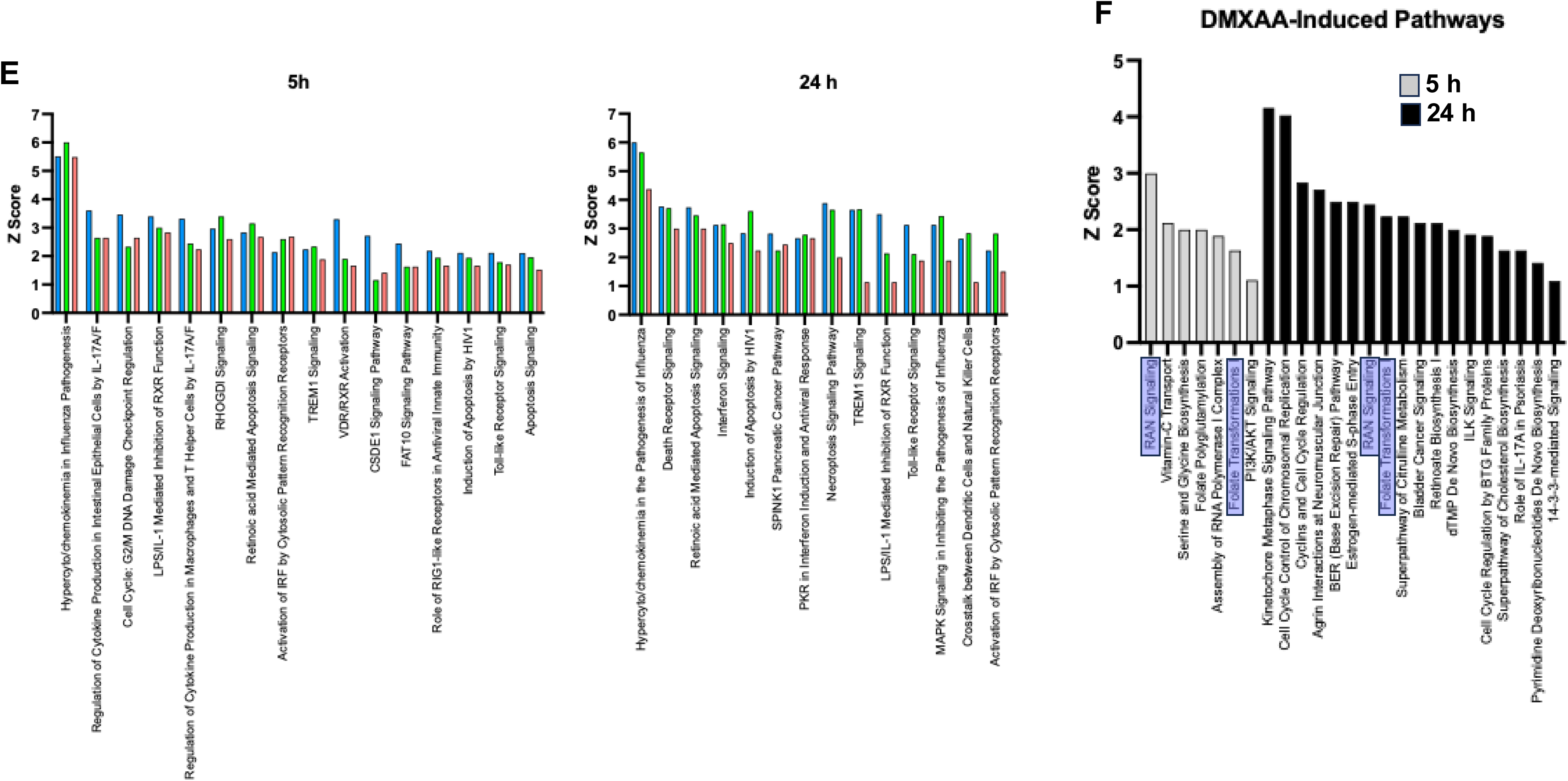
STING agonist-induced transcriptomic changes *in vivo*. C57Bl/6 mice were injected intramuscularly with PBS, CDA, diABZI or DMXAA and total RNA from draining lymph nodes harvested at indicated time point. **A.** Levels of indicated mRNA transcript in draining lymph nodes at 5 h, 24 h, or 72 h for indicated treatment as determined by qPCR. Data presented are mean ± SEM fold changes relative to PBS treated control animals. Statistical significance was determined using a mixed effects ANOVA with Tukey’s correction for multiple comparisons of log transformed fold changes between stimuli at each time point (*p < 0.05; **p < 0.01); **B.** Number of mRNAs significantly up or downregulated more than twofold relative to PBS treatment following indicated treatment as determined by hybridization array; **C.** Principal component analysis of transcript fold changes for indicated stimulus and time point; **D.** Venn diagram of transcripts significantly upregulated for indicated stimulus at 5 h and 24 h; **E.** Signaling pathways displaying a predicted Z score > 1 for all three stimuli at 5 h and 24 h as determined by Ingenuity Pathway Analysis; **F.** Pathways displaying a predicted Z score > 1 following only DMXAA treatment at 5 h and 24 h.

Based on these results we decided to obtain a global picture of changes occurring in whole dLN transcriptomes in response to IM injection of the three agonists. For this we undertook a high content mRNA hybridization array experiment encompassing >21,000 mouse transcripts. Given the degree of temporal diminishment in transcriptional induction (**Figure 4A**), we examined only 5 and 24 h time points. The number of mRNAs showing significant up- or downregulation in dLN harvested following agonist injection were determined by comparing absolute signal levels of mRNA-specific probe binding between STING agonist and PBS injected mice to obtain transcript fold change (**Supplemental Table 1**). As indicated in **Figure 4B**, DMXAA triggers a stronger transcriptional response relative to the other molecules as indicated by the number of significantly induced and repressed mRNAs at both time points. This is also consistent with what is observed by qPCR of targeted genes (**Figure 4A**). Sample clusters obtained using principal component analysis (PCA) further indicated that the transcriptomes generated in response to DMXAA at both time points were substantially more divergent than for the other two stimuli (**Figure 4C**). Consistent with this, at 5 h, stimulation with CDA and diABZI led to upregulation of fewer shared transcripts (779) than were uniquely induced by DMXAA (1642). However, at 24 h diABZI activated considerably more transcripts than did CDA, a larger portion of which were uniquely shared with DMXAA. Over both time points CDA displayed the response of the smallest magnitude and the fewest uniquely induced genes (**Figure 4D**).

We next used pathway analysis to complement quantitative methods and obtain a context-oriented understanding of the biological activities potentially affected by STING agonist administration (**Supplemental Table 2**). We observed within these results pathways similarly activated by all three agonists as shown in **Figure 4E**. Intriguingly, pathways were also detected that were only induced by DMXAA at both 5 h and 24 h (**Figure 4F**). This includes RAN (Ras-related nuclear protein) signaling and Folate transformation pathways, which were predicted to be uniquely induced after both 5 h and 24 h DMXAA treatment. RAN signaling is known to confer inhibitory effects on T cell function largely due to impaired nuclear accumulation of AP-1 transcription factors (c-Jun, c-Fos) [21]. Moreover, folate is known to be important for proliferation of CD8^+^ T cells and may impact CD4^+^/CD8^+^ T cell ratio [22]. These results demonstrate that chemically dissimilar agonists are capable of activating transcription that functionally associates with STING stimulation as well as unique pathways linked with immunologically impactful processes. As such, agonists may confer differential effects on establishment of T cell mediated immune responses.

### Adjuvant-associated changes in cell activation and recruitment to dLN

Innate immune activating adjuvant administration leads to kinetic, activation, and migratory changes in cellular populations at the site of injection and in the dLN [23–26]. This is especially true of APCs such as DC that are essential for initiating antigen-specific T and B cell activation and subsequent differentiation. Importantly, STING activation has been shown to inhibit growth and even induce death of T, B, and monocytic cells [27–30]. How this occurs and its impacts in the context of transient pharmacologic STING activation during vaccination is not known. Given the strong transcriptional response observed in dLN, we used flow cytometry to examine changes in cell populations here 24 h after intramuscular administration of the antigen ovalbumin (OVA) in the presence or absence of CDA, diABZI, and DMXAA using duplicate experiments (**Supplemental Figure 1**). Alum was used as a STING- and IRF3-independent control treatment. As shown in **Figure 5A**, total dLN cell numbers were not significantly altered in response to administration of any stimulus. However, CDA and diABZI led to significant increases and DMXAA significant decreases in B (CD3^-^, CD19^+^) cell composition of dLN while Alum had no effect (**Figure 5B**). DMXAA also led to a significant decrease in the percentage of T (CD3^+^, CD19^-^) cells in the dLN while no other stimulus had an effect (**Figure 5C**). Interestingly, when we examined DC populations, DMXAA led to significant increases in frequencies of cDC1 (CD3/B220/CD11b^-^, CD11c/CD8^+^), cDC2 (CD3/B220^-^, CD11c/CD11b^+^), and pDC (CD3/CD11b/CD8^-^, B220/CD11c^+^) whereas no such changes were observed for the other agonists as shown in **Figure 5C-E**. In addition, DMXAA was also uniquely associated with increases in dLN-localized neutrophils (CD3/B220/F4/80/CD11c-, CD11b^+^/Ly6G^+^; **Figure 5D**) and macrophages (CD3/B220/CD11c^-^, F4/80/CD11b^+^; **Figure 5D**). To understand the mechanistic basis of these unusual DMXAA-mediated observations we asked whether IFN-I responses are required. For this we used mice lacking the IFNα receptor (IFNAR^-/-^). WT and IFNAR^-/-^ mice were treated intramuscularly with OVA alone or admixed with DMXAA. As shown in **Figure 5C**, functional IFN-I signaling was required for increased DMXAA-dependent dLN trafficking of cDC1, cDC2, and pDC but not neutrophils or macrophages. Moreover, IFN-I signaling was also required for DMXAA-dependent decreases in dLN levels of B and T cells. Based on this we conclude that DMXAA induces changes in dLN cell population that are both dependent on and independent of IFN-I signaling. Given the unique transcriptomic and cytokine profiles observed in response to DMXAA it is probable that other factors are also highly relevant and this will require additional investigation.

**Figure 5.**
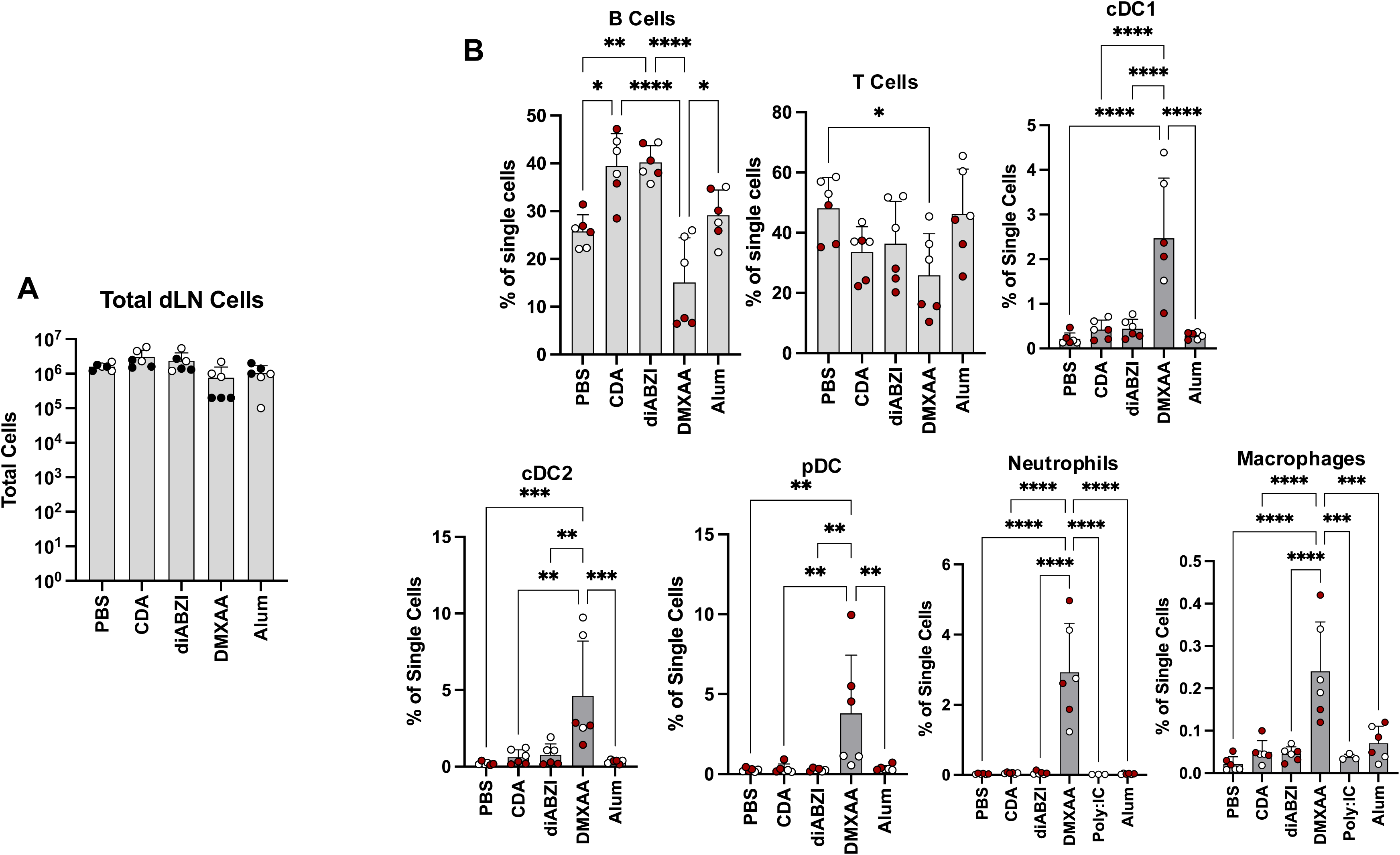
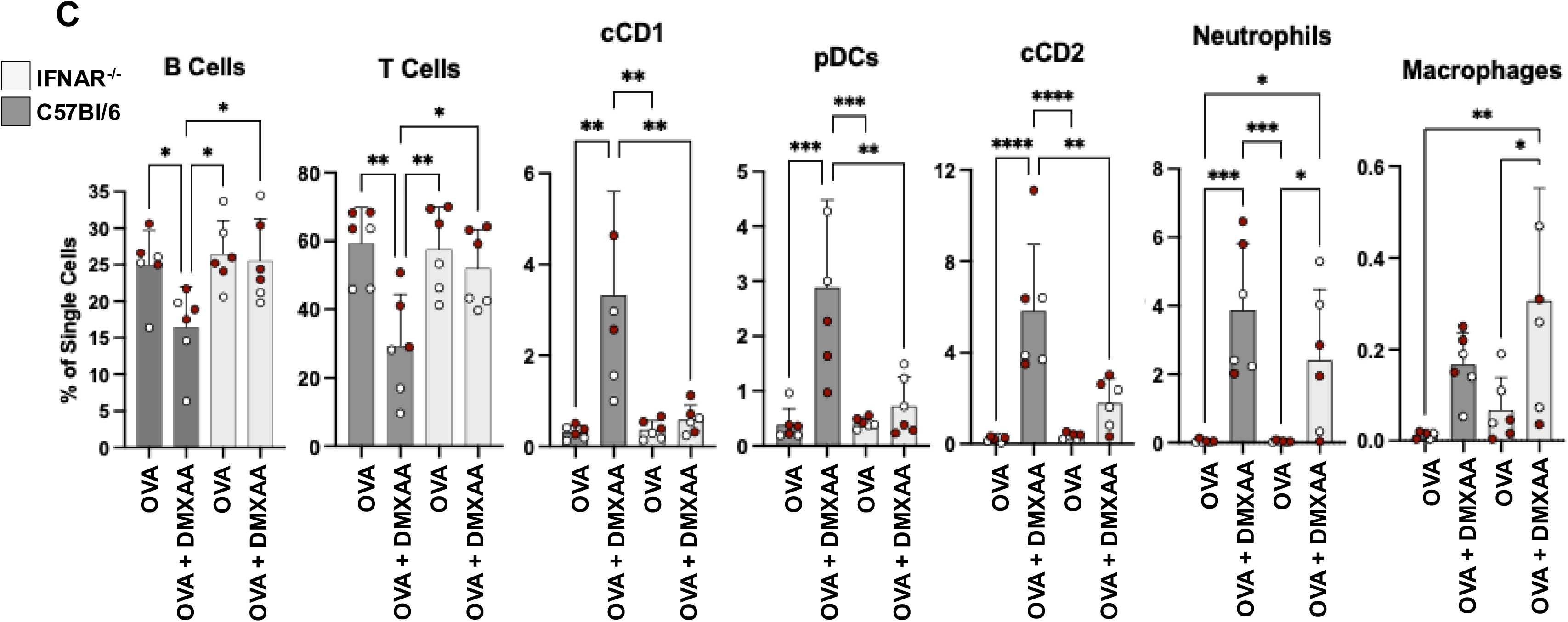
STING agonist-induced changes in draining lymph node immune cell populations. In duplicate experiments C57Bl/6 mice were injected intramuscularly with OVA in the presence of PBS, CDA, diABZI DMXAA, or Alum and single cell suspensions of draining lymph nodes obtained at 24 h post treatment. Flow cytometry was then used to quantify indicated cell types. **A.** Total number of lymph node cells following indicated treatment; **B.** Percent of total lymph node cells represented by indicated B cells, T cells, cDC1, cDC2, pDC, neutrophils, and macrophages as indicated; **C.** Percent of total lymph node cells represented by indicated B cells, T cells, cDC1, cDC2, pDC, neutrophils, and macrophages in WT and IFNAR^-/-^ mice as indicated. Statistical significance was determined using ANOVA with Dunnet’s correction for multiple comparisons (*p < 0.05; **p < 0.01; ***p < 0.001; ****p < 0.0001). Individual experiments indicated by circle color.

### STING ligand-associated enhancement of humoral responses

Given the obvious innate induction stimulated by STING ligands we next compared the extent to which they augment elicitation of antibody responses when co-administered with protein antigen. For this we used a prime/boost strategy via intramuscular injection of OVA. As shown in **Figure 6A**, all ligands led to titers of OVA-reactive total IgG that were significantly higher than observed when OVA was injected in the absence of adjuvant. Additionally, these levels did not significantly differ from those observed when Alum was used. We next examined levels of IgG1 and IgG2c subtypes as an indicator of class switching and T helper (Th) polarization. As shown in **Figures 6B and 6C**, while IgG1 levels for each adjuvant followed the same pattern as total IgG, levels of IgG2c were significantly higher for the STING ligands relative to Alum. This is consistent with what is known about strong Th2 biased immune phenotypes observed for Alum [31] and Th1 biased responses observed for CDA [32]. These results additionally indicate that the small molecule STING ligands diABZI and DMXAA generate T helper polarization that more closely resembles CDA than Alum. Surprisingly, levels of IgG1 but not IgG2c induced by diABZI were significantly lower than those induced by the other STING ligands. This may indicate differential induction of immune processes by the molecule that are mediated by STING or other unknown cellular targets. Regulation of antibody production and affinity maturation is mediated in large part through the function of T follicular helper (T_FH_) cells whose activity can be enhanced by the use of adjuvants [33]. We therefore asked whether adjuvant-specific differences are evident in T_FH_ activation measured following *ex vivo* stimulation of splenocytes with OVA-inclusive peptides. As shown in **Figure 6C**, DMXAA but not the other agonists was able to significantly increase OVA-specific T_FH_ activity following vaccination (**Supplemental Figure 2**). Whether this leads to measurable differences in generation of antibody avidity will require additional studies.

**Figure 6.**
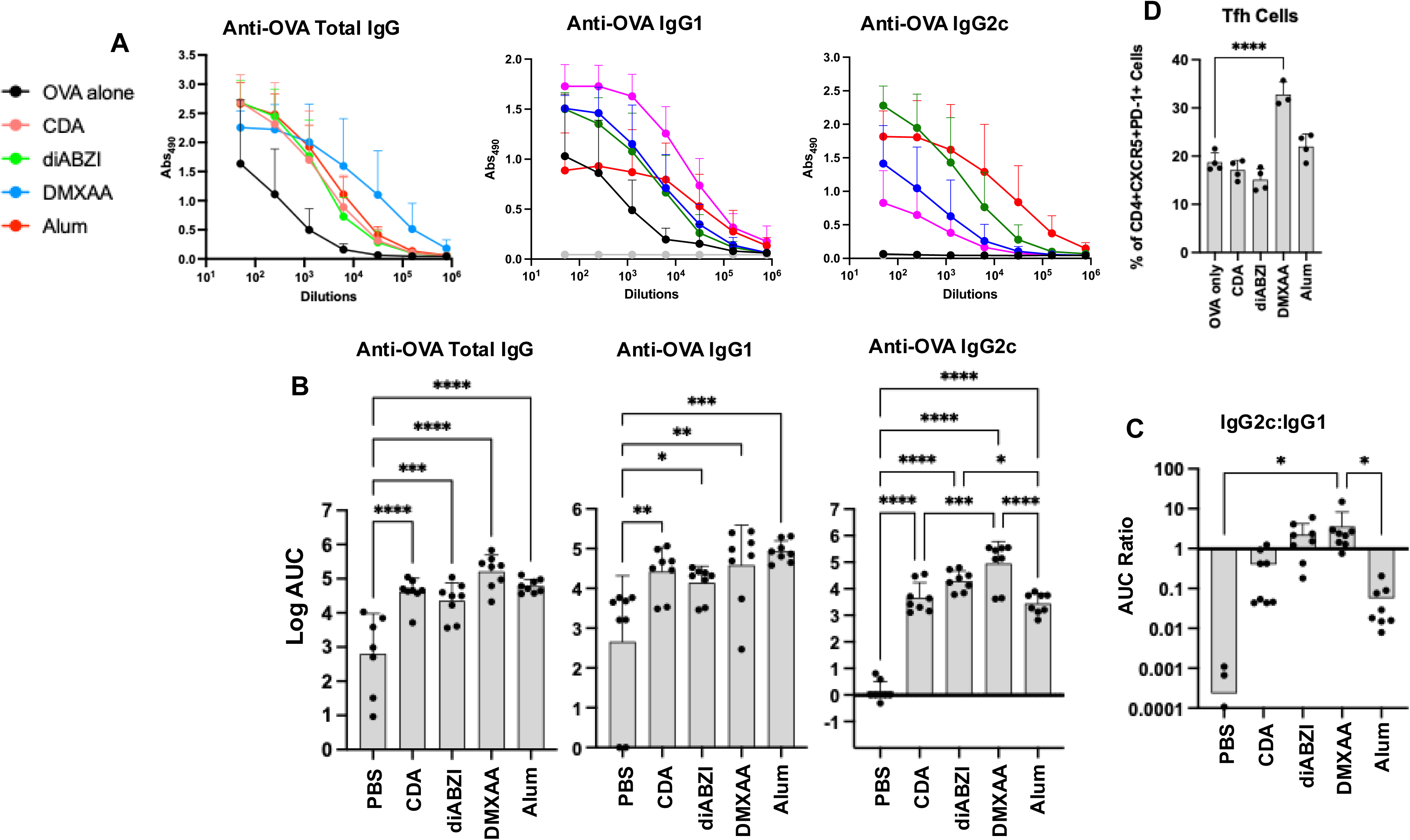
Enhancement of humoral immune response to OVA by co-administered STING agonists. C57Bl/6 mice were vaccinated IM with OVA in the presence of PBS, CDA, diABZI, DMXAA, or Alum, identically boosted 2 w later and harvested 2 w later. **A.** ELISA was performed using serial dilution of harvested sera to quantify levels of total IgG, IgG1, and IgG2c reactive to OVA antigen. Data presented are mean + SEM absorbance (top) as well as mean + SD area under curve (AUC) of absorbance signal including individual animal measurements (bottom); Total number of lymph node cells following indicated treatment; **B.** Mean + SD ratio of IgG2c to IgG1 AUC titers; **C**. Mean + SD percentage of OVA-reactive (OX40/CD25^+^) T_FH_ (CXCR5^+^PD-1high) cells harvested seven days after boost with OVA + indicated adjuvant. Statistical significance was determined using ANOVA with Dunnet’s correction for multiple comparisons (*p < 0.05; **p < 0.01; ***p < 0.001; ****p < 0.0001).

### STING ligand-associated enhancement of cell mediated responses

Previous vaccination models have shown enhanced antigen-directed T cell activation when STING agonists are used as adjuvants [32,34,35]. We therefore used IFNγ ELISPOT to examine whether any differences are observed between the ligands with respect to OVA-directed T cell activation. In duplicate experiments animals were vaccinated as described above and harvested seven days after boost. Splenocytes were then stimulated *ex vivo* using either an OVA-inclusive peptide pool or the MHC class I immunodominant OVA peptide SIINFEKL. As shown in **Figure 7A**, stimulation with the peptide pool led to significant expression of IFNγ in splenocytes harvested from animals adjuvanted with CDA or DMXAA relative to naïve animals or those vaccinated with OVA + PBS. Moreover, CDA adjuvant also led to significant IFNγ expression relative to all other adjuvants following stimulation with SIINFEKL. Expectedly Alum was also unable to elicit any observed IFNγ expression. We next used flow cytometry of splenocytes stimulated with OVA peptide pools to distinguish polyfunctional activation in CD4^+^ and CD8^+^ T cells as indicated by expression levels of IFNγ and TNFα measured following intracellular cytokine staining (ICS) (**Supplemental Figure 3**). As shown in **Figure 7B**, significant expression of TNFα and IFNγ was observed in CD4^+^ and CD8^+^ T cells only following vaccination with CDA. These results suggest that chemical structure of STING ligand associates with differential enhancement of antigen-directed T cell activation.

**Figure 7.**
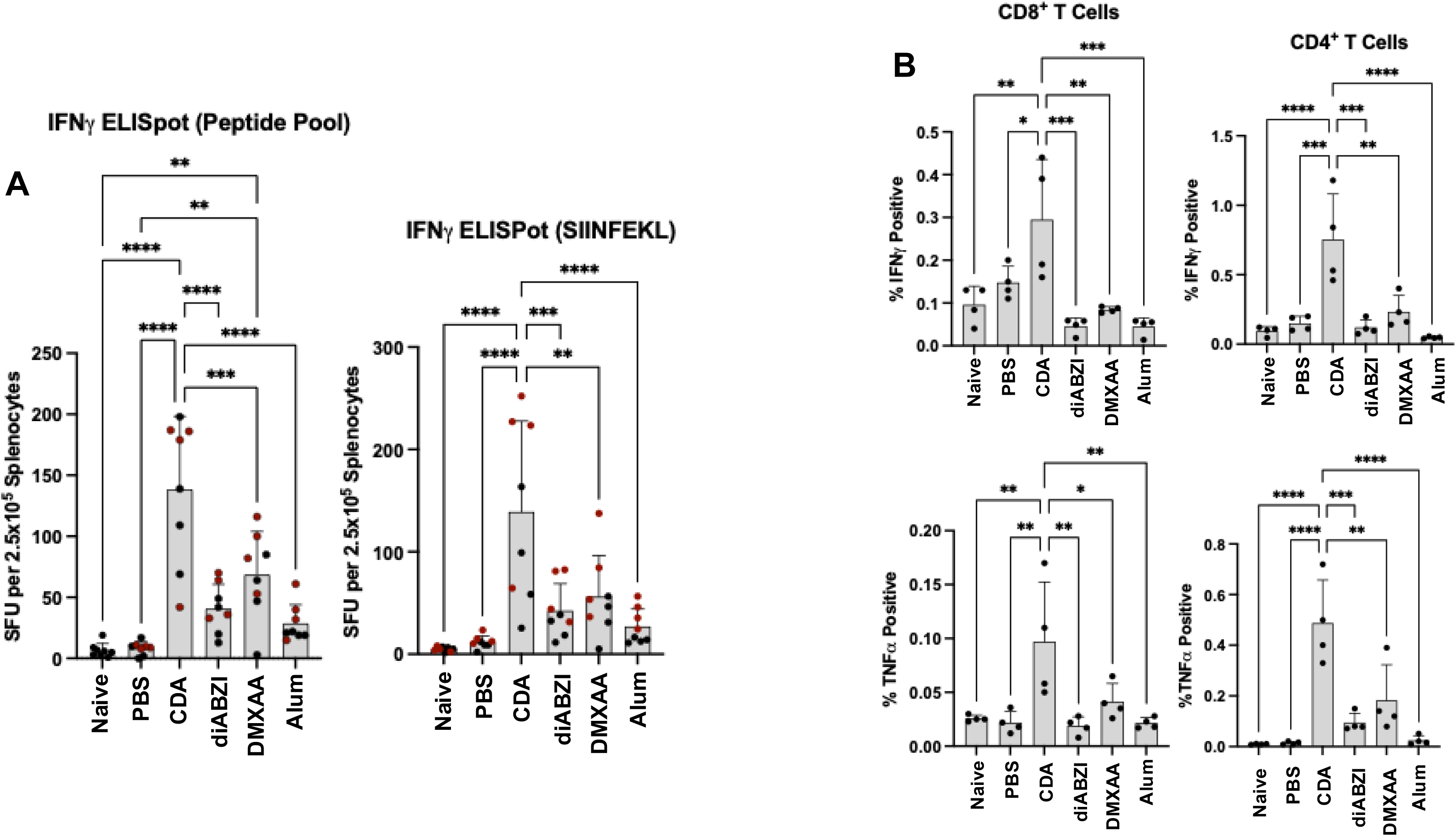
STING-adjuvant associated enhancement of cell mediated responses. C57Bl/6 mice were left untreated or vaccinated IM using a prime/boost strategy with OVA in the presence of PBS, CDA, diABZI, DMXAA, or Alum. **A.** Splenocytes were harvested at 7 d post boost and stimulated *ex vivo* with a peptide pool spanning the OVA protein (left) or SIINFEKL peptide (right). ELISpot was then used to quantify the number of IFNg positive cells. Data presented are mean ± SD number of IFNg positive spot forming units (SFU) per 2.5 x 10^5^ total cells. Results from individual animals are also presented with duplicate experiments represented by marker color; **B.** Intracellular cytokine staining and flow cytometry was used to quantify OVA peptide pool-stimulated expression of IFNg and TNFa from CD4+ and CD8+ T cells in splenocytes. Data presented are mean ± SD percentage of CD8^+^ (left) or CD4^+^ (right) T cells that display positivity for indicated cytokine. Individual animals represented by circles. One way ANOVA using Tukey’s multiple comparison was used to determine statistical significance (*p < 0.05; **p < 0.01; ***p < 0.001; ****p < 0.0001).

## DISCUSSION

Transient pharmacological activation of STING has been shown in animal models to confer desirable immune-mediated outcomes for numerous, seemingly unrelated clinical conditions. This includes enhancement of protective responses conferred by vaccines against viral [35–38], bacterial [39–41], and protozoal [34,42] pathogens. STING agonists have also been shown to impair acute replication of diverse virus types *in vivo* when administered before challenge [43–47]. Moreover, a wide variety of STING agonist platforms have shown remarkable efficacy in mediating T (reviewed in [48]) and NK [49,50] cell-mediated clearance of transplanted tumors including the generation of abscopal responses [9,51,52]. Intriguingly, pharmacologic STING activation has also been shown to alleviate pain (nociception) via IFN-mediated effects on sensory neurons [53] and to attenuate experimental autoimmune encephalitis [54,55]. As such, STING-dependent phenotypes associate with cellular and immunological processes that can be harnessed for diverse clinical benefits and thus pursuit of new agonists of the protein is highly incentivized.

While largely under investigated, the use of exogenous ligands to activate STING can lead to observable differences in phenotypic responses that are linked to chemical structures and protein binding properties of the agonists. The most overt demonstration of this is exemplified by the human-selective agonist C53, which induces activation of STING by engaging the protein’s transmembrane domain [56]. When compared with molecules such as diABZI or cGAMP that activate via the cytosolic domain, C53 is unable (and indeed blocks) induction of TBK1-independent STING-mediated processes such as NLRP3 inflammasome activation, autophagy, and lysosomal biogenesis [7,8,57]. The collective effects of differential induction of these cellular activities on STING-mediated immunological processes are virtually unknown. Additionally, while cyclic dinucleotides including cGAMP, cyclic-di-AMP (CDA) and cyclic di-GMP (CDG) bind to and activate STING, they can also distinctively engage STING-independent PRRs including RECON and ERAdP (CDA [58,59]), or TPL2, DDX41, and TLR7 (CDG [60–62]). These proteins are known to elicit innate immune effects that can enhance or counteract canonical STING-mediated molecular processes through transcription factors such as IRF3 or NF-κB or even induce STING-independent activities such as CREB-dependent transcription. Whether the various classes of small molecule agonists such as those examined here are also capable of activating STING-independent PRRs is unknown. *In vivo*, STING agonist types display differences with respect to cytokine secretion [35,63], enhancement of humoral immunity to subunit vaccines [35,64,65], and cell-mediated tumor clearance [9]. However, an understanding of the combined mechanistic impact of differential STING engagement or induction of other PRRs by STING agonists on downstream *in vivo* immune effects is lacking. As such, studies that comparatively characterize these are necessary to inform future work whose aim is to apply a pharmacologic STING agonism approach toward achieving specific clinical goals.

Work described here demonstrates measurable differences in responses to chemically different STING agonists using *in vitro, ex vivo*, and *in vivo* models. This includes profound differences in transcriptional activity, cytokine secretion profiles, expression of DC maturation markers, elicitation of antigen-directed humoral responses, and cell mediated immune responses. Pharmacokinetic variability is certain to exist between the agonists but given the differences in observed effects *in vitro* and *ex vivo* are unlikely to be solely responsible for the differences detected *in vivo*. This could be due to engagement of STING-independent PRRs, differential induction of STING-mediated processes linked to ligand-induced conformational changes, specific regions of STING binding, or regulation of unknown signaling pathways. These results clearly show that DMXAA generates the most unusual molecular responses despite not displaying atypical relative potency. Interestingly, DMXAA is the only agonist examined here that is species selective, being inactive against primate STING orthologs [12,66]. As such it may engage the protein with unique affinity or kinetics that influence activation properties. Since mammalian STING evolves rapidly under positive selection, likely imposed by the fitness impacts of microbial pathogens [67], species selective agonists are common [13,47,56,68–70]. Moreover, cross-reactive agonists may elicit ortholog-specific molecular activities and thus their use necessitates confirming activation of immunologically relevant phenotypes in both species.

These results also suggest agonist-associated differences in the ability to augment antigen-directed CD4^+^ and CD8^+^ T cell responses with CDA displaying more efficacy in this regard than the small molecule agonists (**Figure 7**). This has relevance for both vaccine enhancement and anti-cancer activity as T cells play key roles in both pathogen clearance and tumor cell killing. Importantly, CDA was observed to induce T cell polyfunctionality as indicated by their expression of both IFNγ and TNFα. As such, better protective immune effects may be demonstrable when using cyclic dinucleotides to potentiate antigen-directed immune responses. Whether this can be mechanistically linked to differential molecular processes induced (or unaffected) by CDA such as activation of NF-κB-dependent transcription (**Figure 1C**), inflammasome activation (**Figure 1D**), or transcriptomic patterns (**Figure 4**) requires additional inquiry but may allow discovery of precise phenotypes desirable for future adjuvant development.

STING represents a valuable pharmacologic target with demonstrated beneficial applications to vaccine enhancement, anti-tumor responses, antiviral protection, and pain suppression. By demonstrating agonist-associated differences in molecular, cellular, and immunological responses this work provides a rationale for pursuing deeper comparative characterization of STING agonists under consideration for clinical use. This includes quantifying cellular processes they induce such as autophagy [6], lysosomal biogenesis [7], and inflammasome activation [5]. Additionally, an understanding cell type-specific differences including agonist sensitivity, transcriptomes, and impact on effector phenotypes will be essential to predict whether the mechanism of action necessary for a desired outcome will be operational across species for a specific agonist. This will also allow identification of criteria essential for validation in both animal and human models that will direct their clinical use.

## MATERIALS AND METHODS

### Reagents and Antibodies

diABZI was purchased from MedChemExpress (Cat # HY-112921B). DMXAA was purchased from ApexBio (Cat # A8233). Sendai virus (SeV) was purchased from Charles River Laboratories (Cat # PI-1). Endotoxin-free ovalbumin was purchased from Invivogen (Cat # vac-pova-100). Steady GLO luciferase reagent was purchased from Promega (Cat # E25100). Quanti-Luc luciferase detection system was purchased from Invivogen (Cat # rep-qlc4lg1). Quanti-Blue SEAP detection reagent was purchased from InVivogen (Cat # rep-qbs). Cell Titer Glo was purchased from Promega (Cat # G7572). Alhydrogel aluminum hydroxide was purchased from Invivogen (Cat # vac-alu-250). ML-RR-S2 CDA was purchased from Invivogen (Cat # tlrl-nacda2r-05). LPS was purchased from Sigma-Aldrich (Cat # L5293). J774-Dual cells were purchased from Invivogen (cat # j774d-nfis). RAW264.7-ISG-Lucia cells were purchased from Invivogen (cat # rawl-isg). GM-CSF and IL-4 were obtained from Peprotech (Cat # 214-14, 315-03. Antibodies used are as follows: anti-mouse CD3 (BioLegend Cat # 100355), anti-mouse CD40 (BioLegend Cat # 124622), anti-mouse CD80 (BioLegend Cat # 124622), anti-mouse CD86 (BioLegend Cat # 105008), anti-mouse CD11c (BioLegend Cat # 117353), anti-mouse CD8α (Thermo Fisher Cat # 45-0081-82), anti-mouse CD19 (BioLegend Cat # 115540), anti-mouse CD45R (BD Bioscience Cat # 553088), anti-mouse CD11b (Thermo Fisher Cat # 48-0112-82), anti-mouse Ly-6G (BD Biosciences Cat # 562700), anti-mouse F4/80 (BioLegend Cat # 123118), anti-mouse IFNγ (BD Biosciences Cat # 554412), anti-mouse TNFα (BioLegend Cat # 506307), anti-mouse CD25 (BD Biosciences Cat #564458), anti-mouse PD-1 (BioLegend Cat #135225), anti-mouse OX40 (BioLegend Cat #119415), anti-mouse Bcl-6 (BD Biosciences Cat # 563363), anti-mouse CD16/CD32 Ab (BD Biosciences Cat #553142), anti-mouse CD45R (BioLegend Cat #103243), anti-mouse CXCR5 (BioLegend Cat #145512), anti-mouse CD3 (BioLegend Cat #100353), anti-mouse CD4 (Thermo Fisher Cat #48-0041-82), and anti-mouse CD8a (Thermo Fisher Cat #MA5-17597).

### Mouse Experiments

All mouse experiments were performed at Oregon Health and Science University in ABSL2 laboratories in compliance with OHSU Institutional Animal Care and Use Committee (IACUC), under protocol 0913. The OHSU IACUC adheres to NIH Office of Laboratory Animal Welfare standards (OLAW welfare assurance A3304-1). C57B6/J or IFNAR^-/-^ mice (aged 5-8 weeks; Jackson Laboratories) were housed in ventilated cage units and cared for under USDA guidelines for laboratory animals. For serum cytokine measurement, transcriptomic, and dLN cell population experiments STING agonists and adjuvant controls were administered intramuscularly (IM) in the quadriceps with DMXAA at 25 mg/kg, diABZI at 1.5 mg/kg, ML-RR-S2 at 0.5 mg/kg, aluminum hydroxide Alum at 15 mg/kg, or PBS. For vaccination experiments animals were primed and identically boosted at 2 w and harvested at 4 w (for antibody readout) or 5 w (for T cell readout). For transcriptomic and draining lymph node experiments STING agonists and other adjuvants were admixed with 10 µg Endofit-ovalbumin (OVA). For T cell proliferation experiments adjuvants were administered with 50 µg Endofit-OVA.

### RAW264.7 and J774 Assays

RAW264.7-ISG-Lucia or J774-Dual cells were seeded to confluence in a 96-well plate overnight in DMEM + 10% FCS. Cells were treated in duplicate with serial dilutions of STING ligands; diABZI and DMXAA starting at 100 µM concentrations. Lipofectamine 3000 (Invitrogen) was used to transfect CDA into RAW264.7 and J774 cells, starting at 100 µM. Lipofectamine was added prior to serial dilutions of CDA. Cells were incubated with STING ligands overnight, 18 h, in 50 uL of DMEM + 2% FBS. Quanti-Luc, Quanti-Blue, or Cell Titer Glo reagents were added (1:1 [v/v]) to each well, and luminescence or absorbance measured on a Synergy plate reader (BioTek). For IL-1β secretion experiments J774 cells were primed 6 h with 100 ng/mL LPS and treated overnight with indicated stimuli. Media was harvested and IL-1β quantified using bead-based Luminex assay per the manufacturer’s instructions (Thermo Fisher Cat # EPX01A-26002-901).

### ELISA Assays

Serological responses specific to OVA were quantified by enzyme-linked immunosorbent assays (ELISA). Blood was collected at 2 w post boost, stored for approx. 1 h at 4° C, and then centrifuged to separate sera. Sera was aspirated and stored at −80 °C until analysis. High-binding microtiter plates were coated with OVA (1 µg/well) (Invivogen) overnight at at 4° C. Coated plates were blocked using 5% (w/v) nonfat dry milk in PBS plus 0.05% Tween 20 (v/v) (PBS-T) for 1 hour at room temperature. Mouse sera was heat-inactivated for 30 mins at 55°C, then serial diluted in 5% nonfat dry milk starting at 1:50 dilution and added to microtiter plate in duplicate. After 2 h incubation plates were washed 3X using accuWash plate washer (Fisher Scientific). Goat anti-mouse IgG-HRP (SouthernBiotech Cat # 1030-05), anti-mouse IgG1-HRP (SouthernBiotech Cat # 1070-05), or anti-mouse IgG2c-HRP (SouthernBiotech Cat # 1079-05) conjugated antibodies (at 1:10,000) were added and incubated for 1 h at room temp. ELISA plates were washed and developed with 0.4 mg/ml o-phenylene diamine buffer (50 nM citric acid, 100 mM dibasic sodium phosphate, pH 5.0). The reaction was halted with 1 N HCl and A_490_ was measured on a BioTek Synergy plate reader. Absorbance was quantified by calculating area under the curve across serial dilutions above baseline detection.

### Cytokine quantification

Proinflammatory cytokines in the sera were examined 5 h after injection. Blood was collected, stored for approx. 1 h at 4°C, and then centrifuged to separate sera which was aliquoted and stored at −80 °C until analysis. Per the manufacturer’s instructions cyto/chemokines from 50 µL sample of each serum aliquot was quantified using a ProcartaPlex Mouse Th1/Th2 multiplex panel (ThermoFisher Cat # EPX260-26088-901).

### RNA isolation and dLN RT-qPCR

Total RNA was isolated from inguinal draining lymph nodes at 5, 24, and 72 hours after IM injection. RNA was treated on column with DNase provided in a Quick RNA miniprep kit (Zymo Research Cat # R1055), according to manufacturer’s protocol. Single-stranded cDNA was generated from total RNA using a RevertAid First Strand cDNA synthesis kit (Thermo Fisher Cat # 1622), using random hexamers to prime first strand synthesis. mRNA expression levels between samples was performed using semiquantitative real-time revere transcription-PCR (qPCR) and quantified by ΔΔC_T_ [71]. Prevalidated Prime-Time 6-carboxyfluorescein qPCR primer/probe sets were obtained from IDT and used for all genes.

### Hybridization Array Analysis

Differential gene expression in lymph nodes of STING agonist IM-injected mice was determined using RNA isolated from the inguinal dLN at 5 and 24 h. Hybridization array data were collected by the OHSU Integrated Genomics Laboratory. RNA sample quantity and purity were measured by UV absorbance at 260, 280, and 230 nm with a NanoDrop 1000 spectrophotometer (ThermoScientific). RNA integrity and size distribution were determined by running total RNA on a Nano chip instrument (Agilent Technologies). RNA was prepared for array hybridization by labeling 100 ng aliquots using the 3’IVT Express kit (Affymetrix). RNA was reverse transcribed to generate first-strand cDNA containing a T7 promoter sequence. A second-strand cDNA synthesis step was performed that converted the single-stranded cDNA into a dsDNA template for transcription. Amplified and biotin-labeled cRNA was generated during the in vitro transcription step. After a magnetic bead purification step to remove enzymes, salts, and unincorporated nucleotides, the cRNA was fragmented. Labeled and fragmented cRNA was combined with hybridization cocktail components and hybridization controls, and 130 µL of each hybridization cocktail containing 6.5 µg of labeled target was injected into a cartridge containing the Clariom S murine array (ThermoFisher cat # 902930) containing >221,900 probes interrogating over 150,000 transcripts from >22,100 genes. Arrays were incubated for 18 h at 45°C, followed by washing and staining on a GeneChip Fluidics Station 450 (Affymetrix) and the associated hybridization wash and stain kit. Arrays were scanned using the GeneChip Scanner 3000 7G with an autoloader. Image inspection was performed manually immediately following each scan. Image processing of sample .DAT files to generate probe intensity .CEL files was performed using the Affymetrix GeneChip Command Console (AGCC) software. Each array file was then analyzed using Transcriptome Analysis Console (TAC; Version 4.0.3) to obtain array performance metrics, calculate transcript fold changes, and perform principal component analysis. To identify probe sets that were significantly regulated in treated versus untreated (mock) cells, we employed a traditional unpaired one-way (single-factor) ANOVA for each pair of condition groups as implemented in TAC. Probe sets were considered differentially regulated if the ANOVA P value was <0.05 and fold change >2. Pathway analysis of significantly regulated mRNAs performed using Ingenuity Pathway Analysis (Qiagen). Venn diagrams were constructed using BioVenn [72].

### Intracellular Cytokine Staining (ICS) assay

Splenocytes from vaccinated mice were incubated with 1 µg/mL OVA peptide pool (#130-099-771; Miltenyi Biotec) at 37°C with 5% CO_2_ for 6 h in the presence of Brefeldin A (#TNB-4506; Tonbo Bioscience) in 96-well U-bottom plates at a concentration of 10^6^ cells/well. Stimulation with 25 ng/mL PMA (Invivogen Cat # tlrl-pma) and 500 ng/mL Ionomycin (Sigma-Aldrich Cat #10634) was included as a positive control. DMSO was used as a negative control. After stimulation, the cells were incubated in anti-mouse CD16/CD32 and stained for 30 min at 4°C with the surface-staining fluorochrome-conjugated Abs: CD3, CD4 (Clone: GK1.5; #48-0041-82; Thermo Fisher Scientific), and CD8a (Clone: CT-CD8a; # MA5-17597; Thermo Fisher Scientific). Intracellular staining was performed for 30 min at 4◦C with the fluorochrome-conjugated anti-mouse IFNγ and anti-mouse TNFα using BD Cytofix/Cytoperm Fixation/Permeabilization kit (BD Biosciences Cat #554714), accordingly to manufacturer’s instructions. Cell events were collected on BD Symphony flow cytometer, and data were analyzed using FlowJo 10 software.

### Activation induced marker (AIM) assay

Splenocytes from vaccinated mice were incubated with 2 µg/mL OVA peptide pool (Miltenyi Cat # #130-099-771) at 37◦C with 5% CO2 for 18h. Stimulation with 5 µg/mL Con A (InVivogen Cat # inh-cona-2) was included as a positive control. After stimulation, cells were stained with LIVE/DEAD Fixable Near-IR Dead Cell Stain Kit (ThermoFisher Scientific Cat # L10119) and blocked with mouse Fc block CD16/CD32 (BD Biosciences Cat #553141). Cells were then stained for 30 min at 4◦C with the following fluorochrome-conjugated anti-mouse Abs: CD3, CD4, CD45R/B220, CD25, PD-1, OX40, and CXCR5. Cell events were collected on LSR-II (BD Biosciences) flow cytometer, and data were analyzed using FlowJo 10 software.

### BMDC activation

Bone marrow extraction was performed by flushing RPMI media through mouse femurs. Cells were pelleted by centrifugation for 5 mins at 500 x g and RBC lysed with ACK lysis (Gibco) buffer at room temp for 3 mins. After centrifugation cells were resuspened in RPMI (1 mL/femur), and counted by trypan blue exclusion (Countess, Invitrogen). Cells were adjusted to 5 x 10^6^ cells per 10 cm petri dish, RPMI was supplemented with 20 ng/mL GM-CSF and 10 ng/ml IL-4 for dendritic cell differentiation. Plates were incubated for 8 d with fresh supplemented media added on days 3 and 6. On day 8 adherent bone marrow derived dendritic cells (BMDC) were chemically lifted from petri dish (Cellstripper, Corning Cat # 25-056-Cl). BMDCs were again counted and plated into 6-well plates at 1-2 x 10^6^ cells/well. Stimuli were added to wells as follows; CDA (1.36 µM), diABZI (17.5 nM), DMXAA (17.5 µM), Alum (3 µg/ml), LPS (100 ng/ml), and DMSO (0.5%). BMDCs were incubated for 20 h, then harvested as before. Antibody staining was performed in FACS buffer [PBS, 0.5% BSA (w/v), 1% 0.5 M EDTA (v/v)]. Fc-mediated binding was blocked with mouse FC-Block (CD16/CD32) (BD Bioscience Cat # 553141), and dead cells stained with Live/Dead Near IR. BMDCs were stained for activation markers CD80, CD86, CD40, and MHC-II for 45 mins on ice. Data acquisition was performed using an LSR-II (BD Biosciences) and analyzed using Flowjo 10.

### APC recruitment

Mice were injected with STING agonists and OVA as described above. Mice euthanized at 24 h time point, and inguinal dLN harvested. Single-cell suspensions were blocked with Fc Block and stained with anti-CD19, anti-B220, anti-CD11b, anti-CD11c, anti-Ly6-G PE-CF594, and anti-F4/80 for 45 mins on ice. Data acquisition was performed using an LSR-II (BD Biosciences) and analyzed using Flowjo 10.

### ELISpot Assays

IFNγ ELISpot assays were conducted as described [73]. Single-cell suspensions of splenocytes were obtained following RBC lysis and added to prewashed mouse IFNγ ELISPot plates (MabTech Cat # 3321-2A). Splenocytes were stimulated with SIINFEKL peptide or OVA peptide pool (10 µg/well), Concanavalin A (0.1 µg/well), PMA/Ion (0.8ng/30ng/well), or 0.5% DMSO. Splenocytes were incubated in ELISpot plates for 18 h. ELISpots were stained according to manufacturer’s protocol. Briefly, plates were washed and incubated with anti-mouse IFNγ biotin antibody for 2 h. Steptavidin-ALP secondary antibody was added for 1 h after plates were washed. Spots were detected using BCIP/NPT-plus substrate, rinsed with water, and dried before counting with an AID ELISpot plate reader.

---

**Supplemental Table 1. STING agonist-mediated dLN transcriptomic changes.** Fold changes of mapped transcripts from hybridization array. Data presented are log2 transformed fold changes for mRNAs and corresponding probe IDs for indicated stimulus relative to mock treated animals and -log(FDRp) values.

**Supplemental Table 2. Stimulus-induced pathway regulation.** Predicted Z scores of indicated biological pathways based on dLN transcriptomic patterns of STING agonists.

**Supplemental Figure 1.** Representative flow cytometry gating plots for dLN B cells, T cells, neutrophils, pDC, cDC1, cDC2, pDC, and macrophages.

**Supplemental Figure 2.** Representative flow cytometry gating plots for IFNγ^+^ and TNFα^+^ CD4^+^ and CD8^+^ T cells.

**Supplemental Figure 3.** Representative flow cytometry gating plots for activated T_FH_ cells.

## Funding

This research was supported by NIH R01 AI143660, R01 AI143660, and HHSN272201400055C.

## CRediT authorship contribution statement

**Victor R. DeFilippis:** Writing – original draft, Validation, Methodology, Formal analysis, Conceptualization, Supervision, Funding acquisition. **Nobuyo Mizuno:** Writing –Methodology, Investigation, Formal analysis. **Dylan Boehm:** Investigation, Formal analysis. **Kevin Jimenez-Perez:** Investigation, Formal analysis. **Jinu Abraham:** Investigation, Formal analysis. **Laura Springgay:** Investigation, Formal analysis. **Ian Rose:** Investigation, Formal analysis.

## Declaration of competing interest

The authors declare no financial interests/personal relationships which may be considered as potential competing interests.

## Notes

### Competing Interest Statement

The authors have declared no competing interest.

