## Supplemental Figure 1 for "Comparative Molecular, Innate, and Adaptive Impacts of Chemically Diverse STING Agonists"

### Slide 1
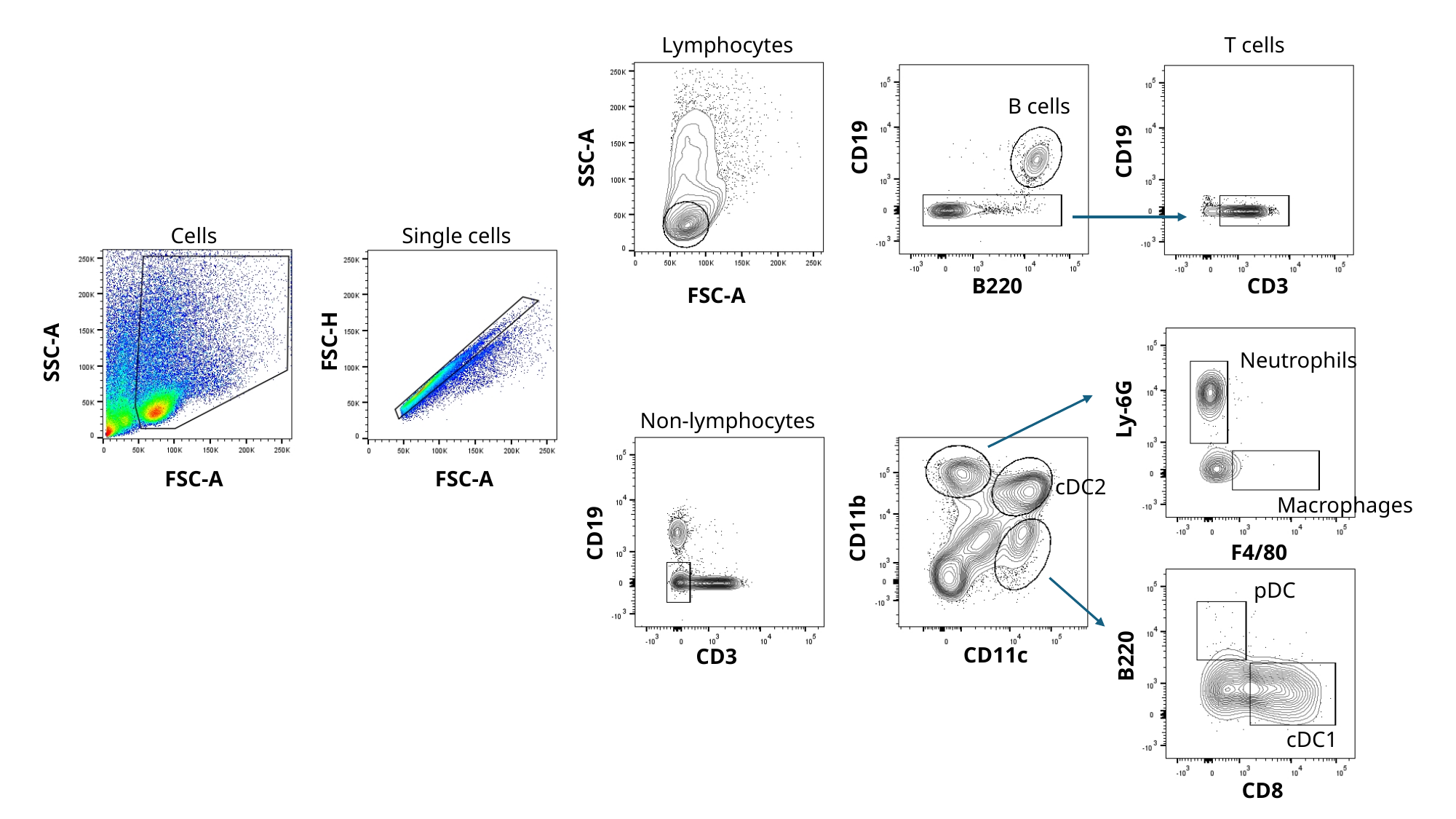

Lymphocytes
T cells
B cells
CD19
CD19
SSC-A
Single cells
Cells
CD3
B220
FSC-A
FSC-H
SSC-A
Neutrophils
Ly-6G
Non-lymphocytes
FSC-A
FSC-A
cDC2
Macrophages
CD11b
CD19
F4/80
pDC
CD11c
B220
CD3
cDC1
CD8
