## Supplementary figures and images for "Comparative Molecular, Innate, and Adaptive Impacts of Chemically Diverse STING Agonists"

### Supplemental Figure 2

#### Slide 1
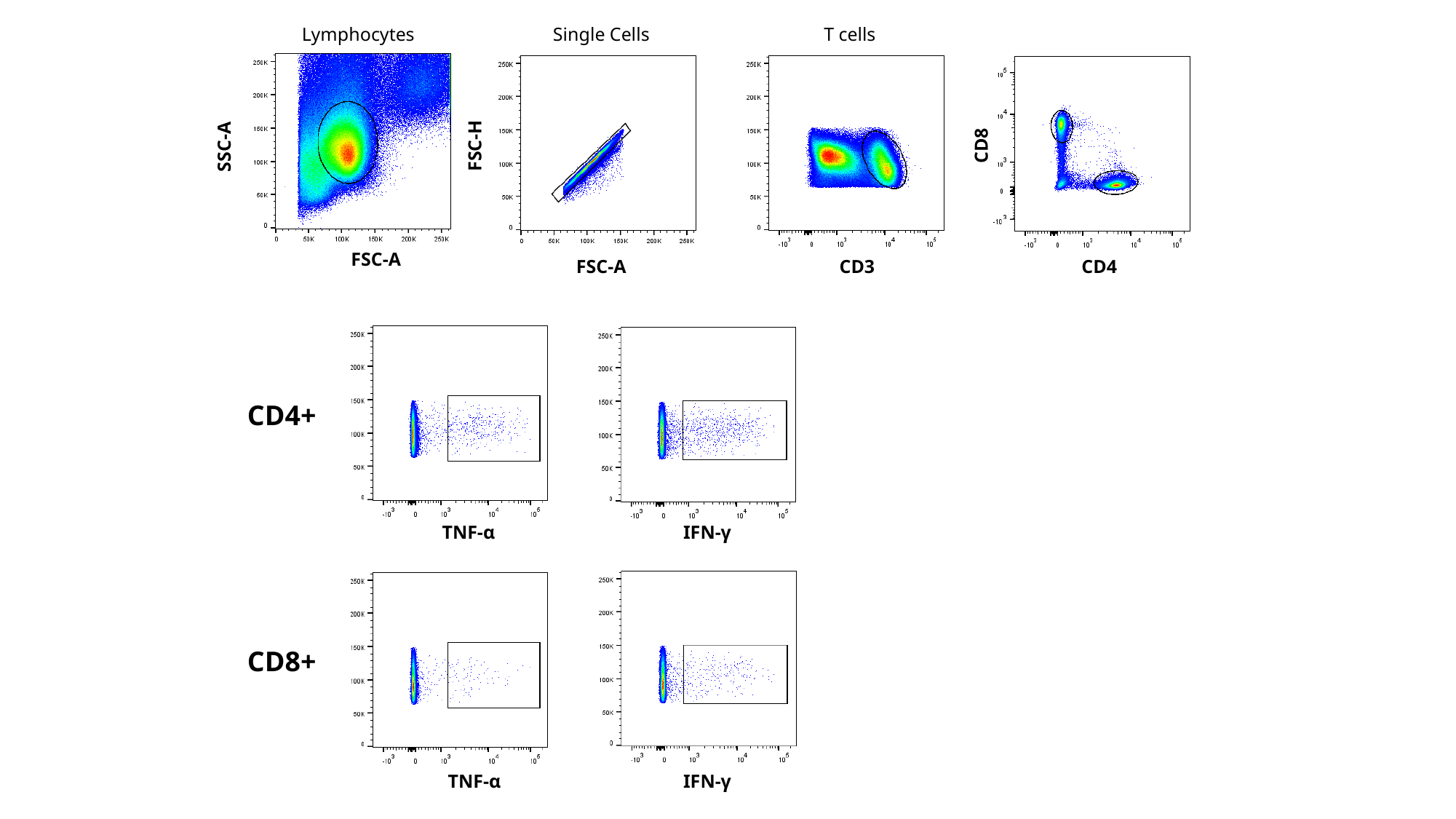

Lymphocytes
Single Cells
T cells
FSC-H
CD8
SSC-A
FSC-A
FSC-A
CD3
CD4
CD4+
TNF-α
IFN-γ
CD8+
TNF-α
IFN-γ

### Supplemental Figure 3

#### Slide 1
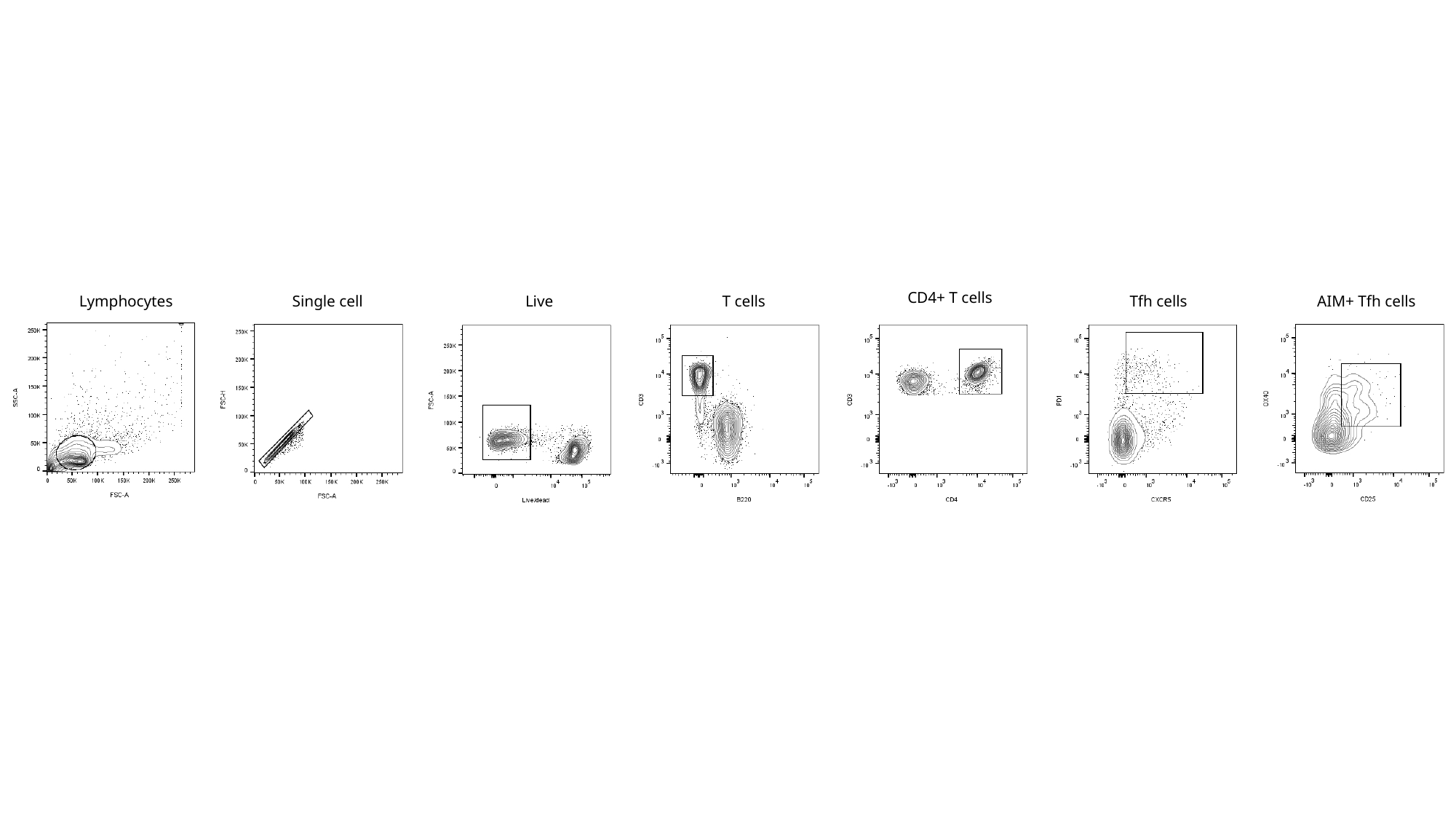

CD4+ T cells
AIM+ Tfh cells
Lymphocytes
Single cell
T cells
Tfh cells
Live
